# Classification of Human Transcription Factors Based on Their Effector Domains via Unsupervised Learning

**DOI:** 10.1101/2025.10.26.684687

**Authors:** Eduardo Ayala, Ayush Gupta, Nehil Shreyash, Arvind Ramanathan, Gül H. Zerze

## Abstract

TFs combine DBDs, which anchor them to DNA, with EDs that regulate transcription through activation or repression, yet the sequence logic linking ED composition to function remains unclear. Here, we systematically define *proxy regions*—disordered segments adjacent to DBDs—to enable quantitative analysis of ED-like sequences across the human TF repertoire. Using a biophysically interpretable 22-feature classifier (FALK22) together with an embedding-based model (ESM), we map ED diversity and identify composition and charge-pattern signatures that correspond to regulatory activity along a disorder continuum, separating activation-from repression-associated regions. FALK22 identified classes align well with those identified from ESM while providing transparent, sequence-level features. Proxy regions near C-termini exhibit gradients that track DBD families, suggesting that EDs and DBDs might have co-evolved rather than evolved independently. These results establish proxy regions and FALK22 as a framework to connect sequence features with transcriptional activity and to generate testable hypotheses about effector-domain function and co-evolution with DNA-binding domains.

**HIGHLIGHTS:**

- We define *proxy regions* as systematically identified disordered segments adjacent to DNA-binding domains (DBD), enabling quantitative analysis of effector domain (ED)-like sequences across the human transcription factor (TF) repertoire.
- We develop FALK22, a 22-feature classification algorithm that classifies transcription factors based on simple sequence properties of their EDs and shows strong alignment with complex embedding-based representations from the Evolutionary Scale Model (ESM).
- FALK22 and ESM uncover distinct amino-acid composition and patterning signatures of EDs that correlate with transcriptional function, separating activation- and repression-associated regions along a disorder continuum.
- Proxy regions located at the C-termini exhibit gradients that correspond to their DBD families, suggesting that EDs did not evolve as independent modular units but rather co-evolved with, or became selectively matched to, their DBD contexts.

## INTRODUCTION

Transcription factors (TF) are key regulatory proteins that control gene expression by binding to specific DNA elements, such as enhancers and promoters, and recruiting the transcriptional machinery necessary to activate a gene^1-5^. Their proper function is essential for maintaining cellular homeostasis, and mutations or dysregulation of TFs (e.g., through overexpression or loss-of-function) are associated with a wide range of diseases^6-8^.

Most TFs contain at least two functionally distinct regions: one or more DNA binding domains (DBD) and effector domains (ED)^9,10^. EDs mediate interactions between TFs and other proteins involved in transcriptional regulation, such as coactivators or corepressors. Although DBDs may be partially or fully disordered before binding, they typically adopt a defined structure upon engaging their target DNA (and binding partners)^11-13^, enabling sequence- and structure-specific interactions. Similarly, EDs—whether functioning as activation or repression domains—often lack stable secondary or tertiary structure in isolation. Some activation domains do acquire defined structures upon binding to their partners^14-16^, but their folding behavior tends to be more variable and highly context-dependent. This structural flexibility supports their ability to interact with a broad range of targets and enables diverse regulatory outcomes. Studies have shown that many transcriptional regulators, including transcription factors, coactivators, mediator complex, and other chromatin remodelers, interact through multivalent, weak interactions (without requiring structure) that drive liquid-liquid phase separation (LLPS), forming dynamic, membrane-less compartments^17-21^.

While TFs are traditionally classified based on their DBDs, whose conserved sequence and recognizable folds enable reliable annotation across species with recognizable folds^11-13,22,23^, ED classification remains underexplored, despite EDs playing critical roles in transcriptional regulation^24,25^. Historically, ED classification has been largely confined to a small number of well-characterized repression domains, such as KRAB, SCAN, and POZ domains, found primarily in specific zinc finger transcription factor families^26-29^. These domain annotations capture a relatively narrow subset of ED diversity and are often insufficient for functional categorization across the full TF repertoire.

More recently, tools for predicting EDs or inferring activation potential have emerged^30-32^, but many rely on datasets derived from yeast^33^, limiting their applicability to humans due to proteomic divergence across species, tissues, and cell types^34-36^. Soto et al.^37^ assembled a compendium of EDs from human TFs based on manual curation of experimental studies. While this resource provides broad coverage and functional annotations, a systematic classification of EDs—and of TFs based on their EDs—using sequence-derived features remained lacking.

To address the lack of sequence-based classification strategies for EDs, here we introduced a 22-feature classification algorithm, Fractions of Amino acids, sequence Length, and Kappa (a measure of sequence charge patterning)^38^, FALK22. Prior studies have linked activation domains to glutamine-, proline-, and acidic-rich regions^39,40^, with recent work demonstrating a correlation between acidic residue content and transcriptional potency^41^. To uncover insights about the role of other amino acids, we incorporated the full amino acid composition into FALK22. Because boundaries of EDs are often ambiguous, we first systematically defined proxy regions (PRs), which are disordered sequence segments adjacent to DBDs, and applied this classifier to PRs, allowing consistent large-scale comparison of ED-like sequences across human TFs.

To complement the FALK22 approach, we also performed a separate classification using embeddings from the Evolutionary Scale Model (ESM), a state-of-the-art protein language model trained on massive sequence databases^42^. These embeddings capture deep contextual relationships between residues but are computationally intensive and less interpretable than FALK22. Both methods were applied to PRs and full-length HTF sequences, enabling a direct comparison between the classification schemes. While the two approaches revealed partially overlapping classes, FALK22, despite its simplicity, achieved clearer segregation of ED types and outperformed ESM in recovering known motif-level distinctions.

## RESULTS

### 0.1 Decomposition of human transcription factors (HTFs) into analyzable sequence subsets

Our search with the keyword “transcription factor”, filtered for human proteins using the most recent UniProt release^43^, identified only 1,465 HTFs. Similarly, previous HTF datasets^34,44,45^ report fewer than 1,600 proteins. The data set constructed by Lambert *et al*.^46^ remains the most comprehensive publicly available resource for HTFs, containing 2,765 entries. We started with this data set and excluded entries marked as lacking “transcription factor activity, sequence-specific DNA binding” as well as those not annotated with “evidence at protein level” (Figure 1f), leaving 1,632 entries. Using the same UniProt release^43^, we then extracted the corresponding protein sequences and annotations. Of these, 802 proteins have only one reported isoform (Figure 1g). For entries with multiple isoforms, we retained only the main isoform, as other isoforms could lack transcriptional activity or DNA-binding capability^47-51^. This procedure defined our “full-sequence” data set.

**Figure 1.**
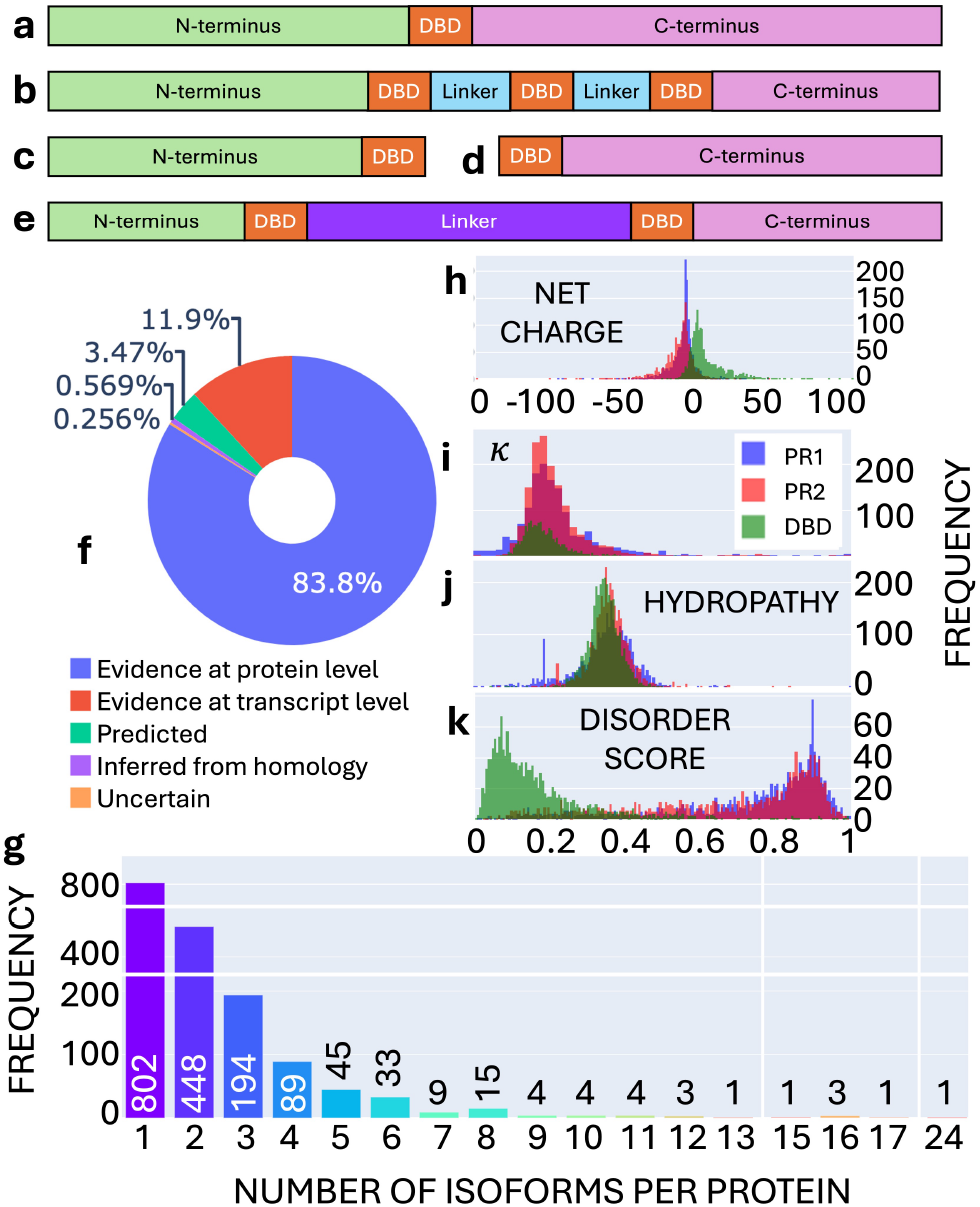
Sequence characterization of HTFs. Examples of the different HTF sequence arrangements: (a) single centrally located DBD with variable N- and C-terminus size, (b) multiple DBDs connected by short sequences identified as linkers. If shorter than 50 amino acids, linkers are not considered as proxy effector domain regions, (c) TFs with a single DBD located at the C-terminus, and (d) N-terminus. (e) A set of 142 TFs contains two DBD regions connected by a long aa segment. (f) Distribution of different existence evidence levels of HTFs. (g) Distribution of the number of isoforms per HTF. (h-k) Distribution of sequence-related metrics for the parameters *κ* (h), net charge (i), normalized hydropathy score (h), and DS (k) for the DBD (*green*), PR1 (*blue*), and PR2 (*red*) regions.

We next analyzed the DBD annotations to approximate ED boundaries and build the corresponding ED datasets. UniProt contains “automatic”, “by similarity”, “ProRule”, and publication-based DBD annotations. We excluded entries without DBD annotations, yielding 1,588 HTFs (with DBD) with at least one annotated DBD. We used these annotations to define the boundaries of the EDs analyzed in our classification scheme. Representative DBD boundary schematics are shown in Figures 1a-e.

Since the precise locations of EDs are often unknown, we identified proxy regions (PRs) as approximations of the regions where activation (or repression) domains could reside. These PRs correspond to extended disordered segments beyond the DBDs, located primarily at the N- or C-terminus (Figures 1a-d) or within long non-DBD linkers (Figure 1e).

For each HTF, our aim was to identify at most 2 PRs. Approximately 57% (954 HTFs) contain more than one DBD (e.g., Figure 1b). When two DBDs are separated by linkers shorter than 50 amino acids (aa), linkers were not considered part of any PR. However, 142 HTFs contain longer linkers (*>* 50 aa) between two DBDs (Figure 1e). Such longer linkers could stabilize chromatin and indirectly regulate transcription by mediating specific protein-protein interactions to bridging between distant enhancer/promoter regions (if permitted by steric hindrance)^37,52^; therefore, we included them as a proxy region. We found that for all TFs, the disordered C-terminus tail (if present) is consistently longer than the any linker segment and hence was designated PR1. Any significantly long linker fragment designated PR2 if it is at least 50 residues longer than the N-terminal tail; otherwise, the N-terminal tail was assigned as PR2. Using these criteria, we compiled the 1631 sequences in the PR1 database (C-termini) and the 1529 sequences in the PR2 database (linkers or N-termini). The compiled full-length, PR1, PR2, and DBD databases are provided as Data S1.

### 0.2 Hydropathy, charge, and disorder profiles of DBDs and PRs

Intrinsically disordered proteins (IDP) and normally foldable (globular) proteins are traditionally separated on a pseudophase diagram proposed by Uversky *et al*.^53,54^, defined by mean net charge (MNC) and mean hydropathy score^55^, commonly known as the Uversky plot (Figures S2a, S2d). However, this classical approach placed 45% and 43% of the PR1 and PR2 sequences, respectively, in the ordered region (Figure S2a and S2c). This can also be seen in Figure 1, j, which shows that the hydropathy distributions for the PRs were centered very close to those for DBDs. The MNC-hydropathy features generate an overly dense space that lacks clear segregation, leading to a poor clustering performance.

As an alternative to the hydropathy index, we calculate the disorder score (DS) introduced by Emenecker et al.^56^, which is a deep-learning predictor trained on consensus annotations from multiple methods to yield a normalized per-residue probability of disorder. It provides a more robust scale for cross-protein comparisons since it integrates diverse sources of experimental and computational information. DS values for PR1 and PR2 are centered around 0.9 (Figure 1k), confirming their predominantly disordered character and placing them and DBDs at the opposite ends of the spectrum. To further contextualize these regions relative to other sequence types, we compared their average sequence properties—including hydropathy, net charge, DS, and individual amino acid fractions—to those of full-length TFs, their DBDs, and the average human proteome (Figure S1). This comparison reveals that DBD are distinctly enriched in positively charged amino acids, cysteine, and tyrosine, whereas PRs are enriched in acidic residues, other polar amino acids, and prolines.

Applying HDBSCAN^57^, a density-based clustering technique, to MNC-hydropathy Uversky space resulted in *>* 50% of proteins classified as noise in every trial (Figures S2b and S2d). A related representation proposed by Das et al.^58^, which plots the fractions of positive (*f* ^+^) and negative (*f* ^−^) residues, produced a similarly dense distribution (Figure S3) and an even higher proportion of noise points. These results confirm that classical two-feature charge—hydropathy spaces cannot effectively separate DBDs and proxy regions. Numerous studies have extended such pseudophase diagrams to include conformational states of IDPs as additional functions of charge patterning^38^. These sequence-based approaches have also been used as guides in both experimental^59,60^ and computational^61^ studies of IDPs.

Since neither of these classical approaches provided adequate sequence differentiation, we next explored whether further features could better distinguish DBDs and PRs. We found that DBDs are slightly positively charged on average (Figure 1h), consistent with predictions of an overall positive charge on DBDs^62^. Furthermore, we found that the EDs are predominantly negatively charged (Figure 1h), which is consistent with the fact that acidic amino acids are common in the activation domains, which are considered to regulate the binding of DBD and DNA (e.g., being electrostatically repulsive)^63^. Beyond the net charge, the spatial arrangement of the charged residues—quantified by the charge patterning parameter *κ*^38^—also serves as a differentiating factor. *κ* ranges from 0 to 1, where low values correspond to well-mixed opposite charges, while values near 1 indicate segregated charge blocks. *κ* ranges from 0 to 1, where low values correspond to well-mixed opposite charges, while values near 1 indicate segregated charge blocks. For both DBDs and proxy regions, the *κ* values are centered around 0.16 (Figure 1h), where PR1 has a broader distribution of *κ* compared to PR2. In FALK22, we also used *κ* as an informative descriptor.

Analysis of PR lengths revealed that 278 of the HTFs have PR1 segments shorter than 10aa, while 66 have PR2 segments below this threshold (Figure S4a). Based on the calculated average DS per residue (Figure S4b), which drops near 10 aa and crosses the disorder-order threshold of 0.5, such short segments are unlikely to form stable motifs^64^ and are too short to engage in extended intermolecular contacts. This observation further motivated the inclusion of sequence length as a key descriptor in FALK22.

Collectively, these analyses prompted the development of a broader yet still minimal feature set, FALK22, for systematic classification as presented in the following section.

### 0.3 Feature representation for PRs as surrogates for EDs

EDs derive their activity from sequence composition and disorder rather than well-defined structure. Building on evidence that activation domains often rely on aromatic, leucine, and acidic residues^30,41,65,66^, we first tested whether similar compositional and charge-based descriptors could distinguish the PRs identified in HTFs.

We began with a seven-feature representation (F7) that captured the minimal physicochemical signatures previously linked to activation and repression domains: mean hydropathy, fractions of isoleucine, proline, and glutamine, fractions of positively (*f* ^+^) and negatively (*f* ^−^) charged residues, and the mean net charge per residue (MNC = |*f* ^+^*f* ^−^ |). UMAP projections (which is a dimensionality reduction technique) of this space revealed no consistent gradients with respect to disorder score (DS), mean hydropathy, or MNC for either PR1 or PR2 (Figures S5a–S5l). Motif-enrichment analyses using FIMO^67^ and STREME^68^ identified dispersed motifs (Figures S5c,f,i,l), consistent with the absence of well-defined subgroups, in contrast to structured motif clusters within the KRAB, SCAN, and BTB/POZ families^27^.

To incorporate additional sequence-level information neglected in F7, we added the charge-patterning parameter *κ*^38^ and sequence length—both we hypothesized to influence conformational and phase-separation behavior—forming a nine-feature representation (F9). The resulting projections (Figures S5m–x) showed only subtle shifts in cluster density, indicating that *κ* and length alone do not significantly improve separation. These findings implied that ED-like regions cannot be characterized solely by a few canonical amino-acid enrichments or by overall charge metrics.

We therefore constructed a more comprehensive 22-feature space, FALK22 (Fractions of Amino acids, sequence Length, and Kappa), encompassing all amino-acid fractions, *κ*, and sequence length. FALK22 captured clear gradients of disorder across PR1 and PR2 (Figures 2a, 2e), (as well as full-length sequence, Figure 2i) with PR2 exhibiting broader DS variability, consistent with its broader DS distribution (Figure 1k). Importantly, FALK22 also resolved distinct family-level organization when colored by DBD identity (Figures 2b, 2f). Among the largest families (Figure 2q), the Nuclear Receptor (NR) and Forkhead families were clearly segregated on PR1 projection (Figure 2b, the coloring of the projection is as bar colors in Figure 2q), whereas C2H2 Zinc-Finger (ZF) and Homeodomain families were clearly segregated on PR2 projection (Figure 2f).

**Figure 2.**
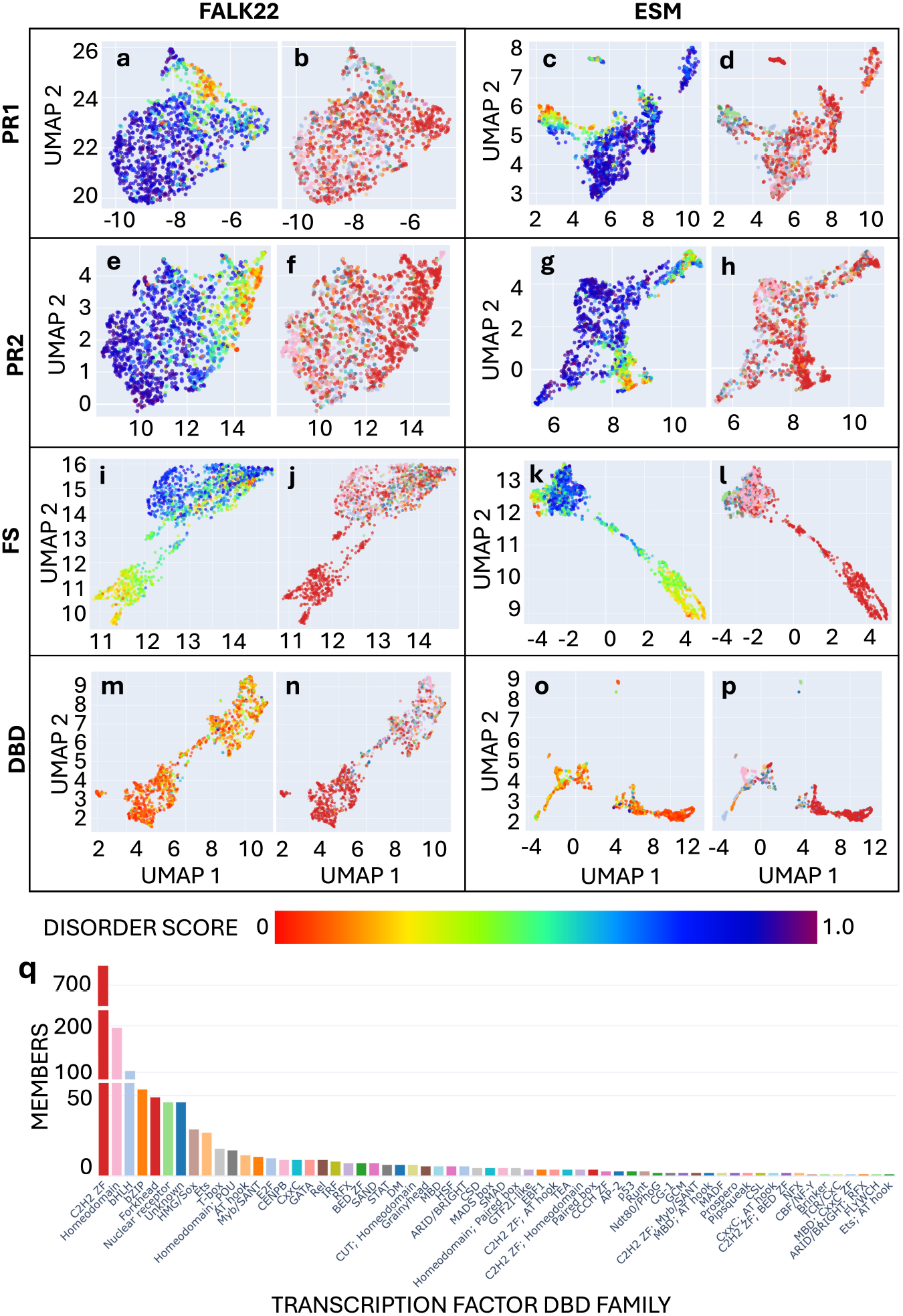
Two-dimensional projections of high dimensional FALK22 and ESM-based features, colored by the two properties that showed strongest gradients. Two-dimensional projections for PR1 and PR2 of FALK22 and ESM colored with respect to (a, c, e, g, i, k, m, o) DS and (b, d, f, h, j, l, n, p) DBD family. The DBD family color scheme for the transcription factors is shown in the size distribution plot (q). This family coloring scheme is kept consistent throughout all figures whereever DBD family based coloring is applied. ESM projections were reduced to 2 dimensions for comparison purposes.

PRs spanned the entire disorder range, consistent with their functional plasticity. NR family EDs clustered toward the more ordered region (low DS region)—matching their structured activation domains known to stabilize DNA–protein interactions (Figures 2a and 2b). Similarly, the C2H2-ZF family populated mostly the low DS region in PR2 projections (Figures 2e and 2f). These gradients and family-specific arrangements demonstrate that FALK22 captures biophysical diversity that mirrors both transcriptional function and DBD-ED evolutionary context.

When we processed the full-length sequences (FS) of HTFs as well as their DBD sequences, Comparable trends were also observed when the same features were applied to the full-length HTF sequences (FS) and their isolated DBD segments, both of which exhibited weaker but same disorder and DBD family gradients (Figures 2i, 2j, 2m, and 2n). These consistent relationships across domain-level and full-protein projections indicate that the physicochemical principles encoded in FALK22 generalize beyond the proxy-region datasets, capturing hierarchical organization from individual domains to entire TFs. The same gradients strengthened going from FALK22 to a context-aware language model (Figures 2k, 2l, 2o, and 2p) as it will be discussed in the next subsection. Gradients on further features will be also discussed in the next subsection (Figure 3).

**Figure 3.**
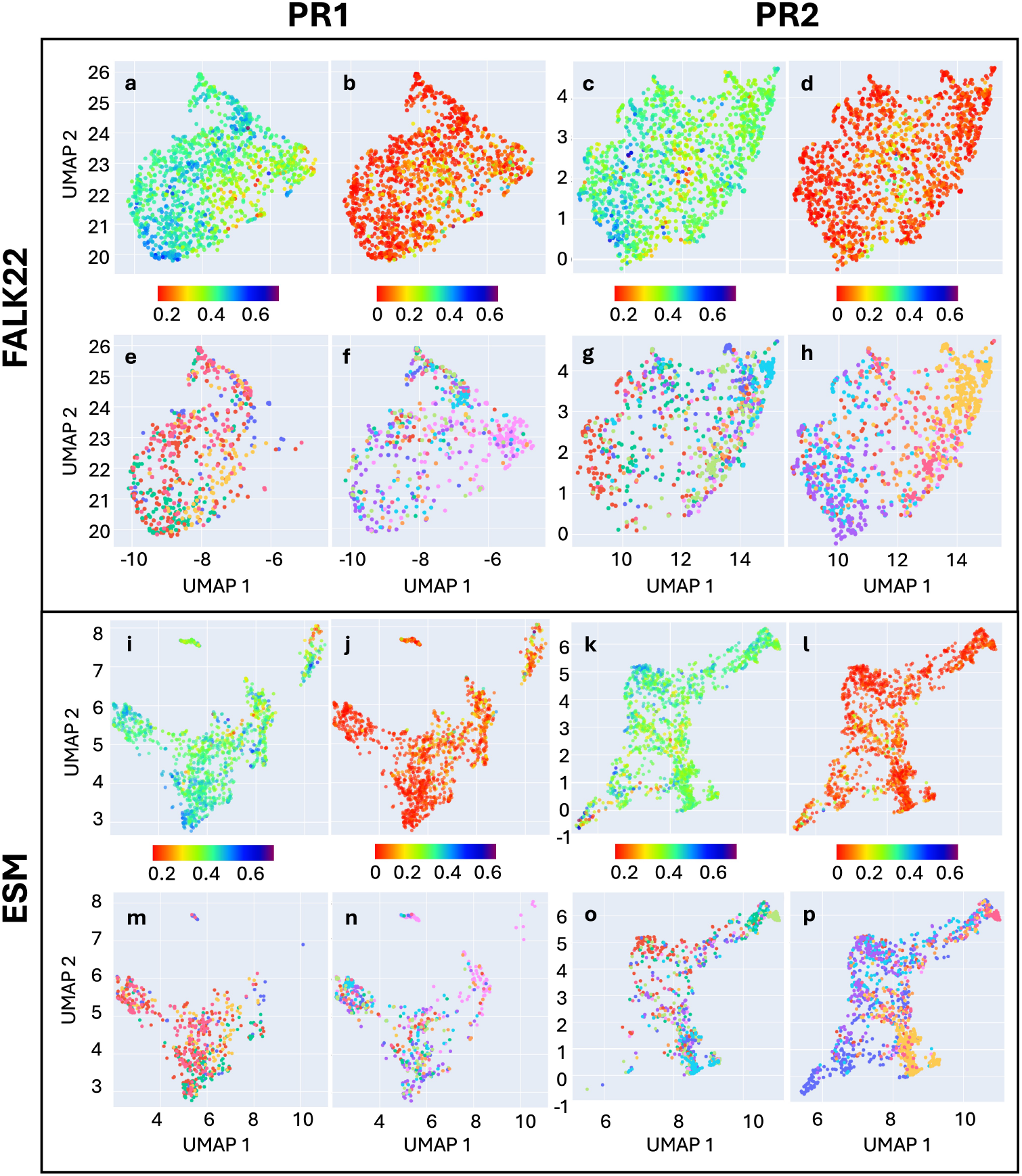
Two-dimensional projections of high dimensional FALK22 and ESM-based features for PR1 and PR2, colored by four other properties. (a-h) Two-dimensional projections for PR1 and PR2 of FALK22 and (i-p) ESM. (a, c, i, k) Colored projections with respect to the hydropathy score, (b, d, j, l) MNC, with the corresponding colorscales below the panels. (e, m, g, o) FIMO and (f, n, h, p) STREME sequence motifs (wherever successfully found) distribution over the sequence space. The color key for the motifs is not shown for simplicity. But the assignment of the same color indicate presence of shared motifs.

While FALK22 provides an interpretable, composition-based description of sequence space, it does not explicitly encode positional context (positions of aa with respect to each other) or higher-order correlations between residues—features that may underlie motif co-occurrence, cooperative binding, and post-translational modification patterns. To evaluate the extent of the importance of contextual information, in the next subsection, we compared FALK22 to a transformer-based, context-aware language model trained on millions of natural protein sequences.

### 0.4 Comparison of FALK22 with a large language model

Context-aware large language model (LLM) embeddings have achieved remarkable success in predicting post-translational modifications (PTMs),^69^ structural similarities,^70^ and more recently, complete 3D structure prediction.^42^ We expect such representations to implicitly inform about the protein’s regulatory partners (ligands, coactivators, repressors, etc.) in the case of HTFs. To benchmark FALK22 against such deep-learning LLM representations, we compared it with token embeddings from the latest version of the Evolutionary Scale Model (ESM-2)^42^, a transformer model trained on ∼65 million UniRef sequences^71^.

For both PR1 and PR2, two-dimensional UMAP projections of ESM embeddings (Figures 2c, 2d, 2g, and 2h) produced denser, less separated spaces than FALK22 but retained the same global gradient in disorder score (DS) (Figures 2a and 2c; as well as 2e and 2g). When colored by DBD family, ESM reproduced the major groupings observed in FALK22—especially the compact nuclear-receptor (NR) cluster in PR1 (Figures 2b and 2d, pale green family) and the dispersed C2H2-ZF family in PR2 (Figures 2f and 2h, dark red family). Strinkingly, these very high-dimensional embeddings generated by an LLM trained over 65 million protein sequences produced the same gradients as the simple 22-descriptor FALK22. This finding suggests that family-specific biophysical trends are compositionally encoded rather than being contextual. We repeated the feature mapping for FS and isolated DBD segments. FS maps showed stronger gradients of both DS and DBD families on reduced dimensionality (UMAP) projections (Figures 2k and 2l) with ESM, where the NR family matched to low DS at the top left corner, C2H2 ZF family also matched to low DS at the bottom right corner, and homeodomain family matched to high DS at the top left corner. Importantly, these consistent relationships across domain-level and full-protein projections reinforce the same conclusion that similar physicochemical information is encoded within these regions, suggesting a possible co-evolution of DBDs and EDs.

Quantitative analysis of family proximity (Figure S6) revealed that within-family distances are smaller for PR1 than PR2 across both models, indicating that C-terminal EDs (PR1) are more conserved in their physicochemical signatures. PR2 sequences, typically at N-termini (Figure S7), showed greater dispersion, consistent with their diverse roles in protein—protein interaction networks. These differences imply that while EDs share no sequence homology, their disorder and charge patterns preserve family-specific physicochemical fingerprints, supporting the notion that EDs co-evolved with DBDs.

In addition to the DS and DBD family gradients, we also examined whether these feature projections show gradients on other properties (Figure 3). In FALK22, PR1 and PR2 embeddings show subtle gradients with respect to hydropathy and MNC—two properties known to influence intrinsic disorder^72^—whereas these gradients are absent in ESM (Figures 3a–d vs 3i–l). The existence of such gradients in FALK22 implies that it captures additional axes of physicochemical variability, potentially linked to differences in phase-separation or condensate-forming propensities^73^. We also tested whether motif-level organization was preserved by mapping FIMO- and STREME-identified motifs onto each feature space. FALK22 revealed dispersed motif distributions for PR1 and more compact, domain-like enrichment for PR2 (Figures 3e–h), whereas ESM embeddings showed weaker motif segregation (Figures 3m–p). These results demonstrate that despite its simplicity, FALK22 can recover both compositional and some contextual organization comparable to that of transformer-based embeddings.

To determine how similar or different the representations from FALK22 and ESM are, we also performed a feature space alignment in three dimensions after normalization of the ESM embeddings and FALK22. We used Umeyama’s algorithm^74^, which uses singular value decomposition to find the optimal rotation matrix and translation vector that minimizes the sum of squared distances between the two spaces. The aligned spaces in Figure 4 show that for both PRs, the FALK22 space largely differs in shape and distribution when compared to the ESM embeddings. The PR1 and PR2 projection alignments have an average RMSD of 0.6025 and 0.3626 (Figures 4b and 4c), respectively. This made us question the capability of ESM to represent the PRs, since the ESM is trained on complete sequences (i.e., not fragments) and contains separate tokens that define the start and end of the sequence, potentially limiting its utility to study complete proteins only. This type of influence may degrade the representation for any of the PRs, as none of those sequences is complete by itself. To address this hypothesis, we also derived the reduced features for the FS dataset using FALK22 and ESM and projected them in 3D space following the alignment procedure defined above (Figure 4a). Strikingly, the aligned representations have a very similar shape, with a smaller RMSD of 0.3152 for the FS dataset.

**Figure 4.**
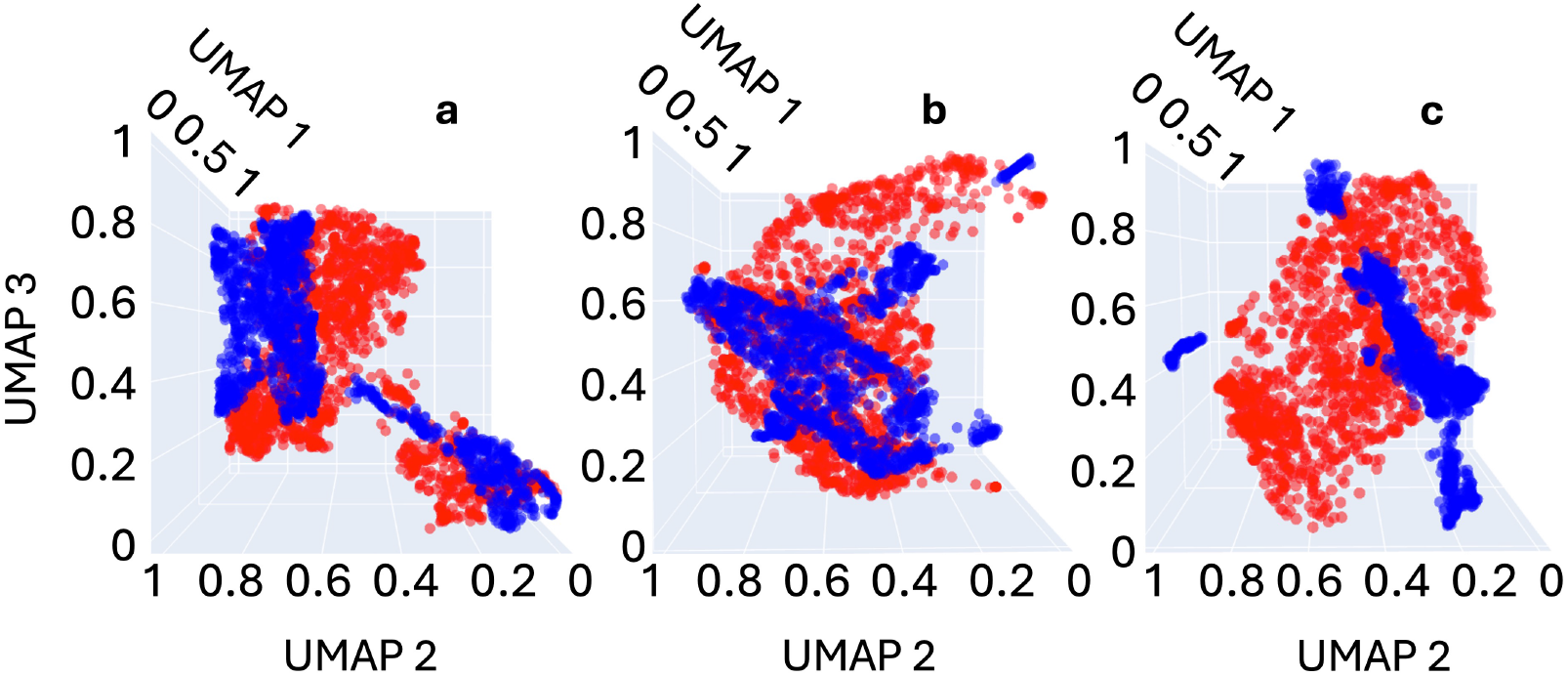
Alignment of FALK22 and ESM embeddings in the three-dimensional space. FALK22 (*red*) and ESM embeddings (*blue*) for a) FS (RMSD: 0.3152), b) PR1 (RMSD: 0.6025) and c) PR2 (RMSD: 0.3626).

We note that we found this large resemblance and the close alignment only for the ESM model that was trained with 150M parameters. Increasing ESM size to 650M or 3B parameters drastically altered embedding geometry and reduced motif segregation (Figure S8, Table S2), echoing known performance saturation effects in large language models^75^. This analysis shows that going above 150M parameters is not only more computationally expensive but also has poorer performance. By contrast, the 22-feature FALK22 model maintains stable geometry, interpretable axes, and minimal computational cost while producing comparable clustering and family-level organization. Having optimized this interpretable feature space, we next applied clustering analyses to classify human TFs according to their effector-domain properties.

### 0.5 ED based classification of HTFs

We next applied HDBSCAN to classify the HTFs based on their FALK22-derived PR features. Parameter optimization (Figure S9) minimized noise and produced seven primary PR1 clusters and twelve primary PR2 clusters initally, which are then further refined via subsequent clustering. After the first round of clustering, the PR1 dataset splits into 7 clusters with moderately low noise of 27% (Figure 5a, left panel). Mutual-information analysis (Figure 5a, right panel) identified sequence length, proline, and alanine fractions as the dominant discriminants for the largest clusters of PR1, with subsequent refinement revealing the influence of methionine and cysteine fractions (Figures 5b to 5e, right panels), as well.

**Figure 5.**
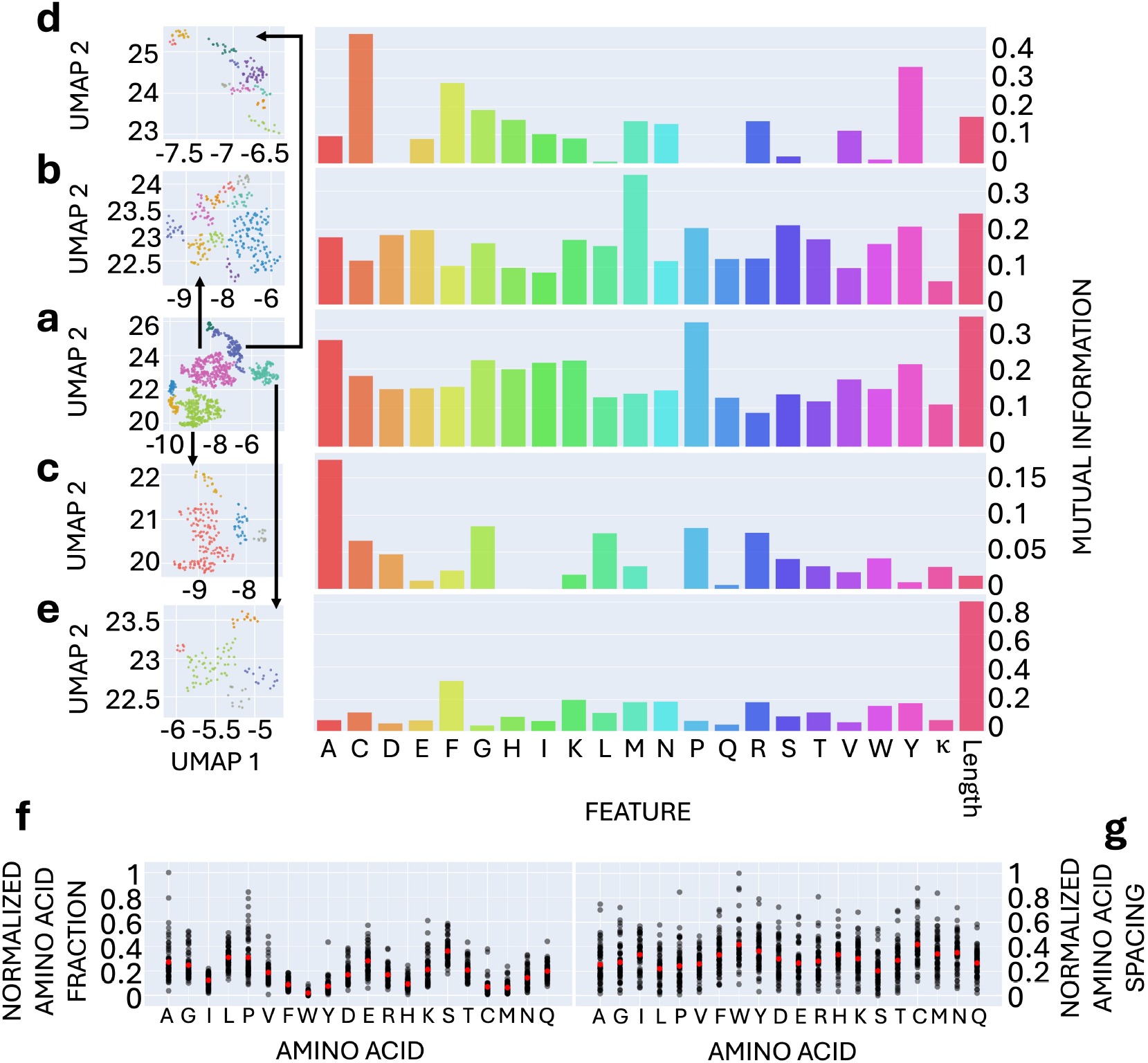
Clustering of the 1,361 elements in PR1 with its corresponding mutual information plot. a) Initial set split into 7 clusters with 36, 40, 48, 114, 152, 295, and 309 sequences. Projections included the subsequent clustering of the 309 (b), 295 (c), 152 (d), and 114 (e). Further clustering steps are excluded for visual clarity. Noise points are not included in the projections. Normalized averages per amino acid fraction (f) and identical amino acid spacing (g) for the obtained clusters (*black*) and global averages (*red*).

PR2 dataset initially splits into 12 clusters with moderate noise of 35% (Figure 6a, left panel). For PR2, alanine, proline, tryptophan, and tyrosine, and sequence length appear to be key differentiators for the generated clusters (Figures 6a to 6d, right panels).

**Figure 6.**
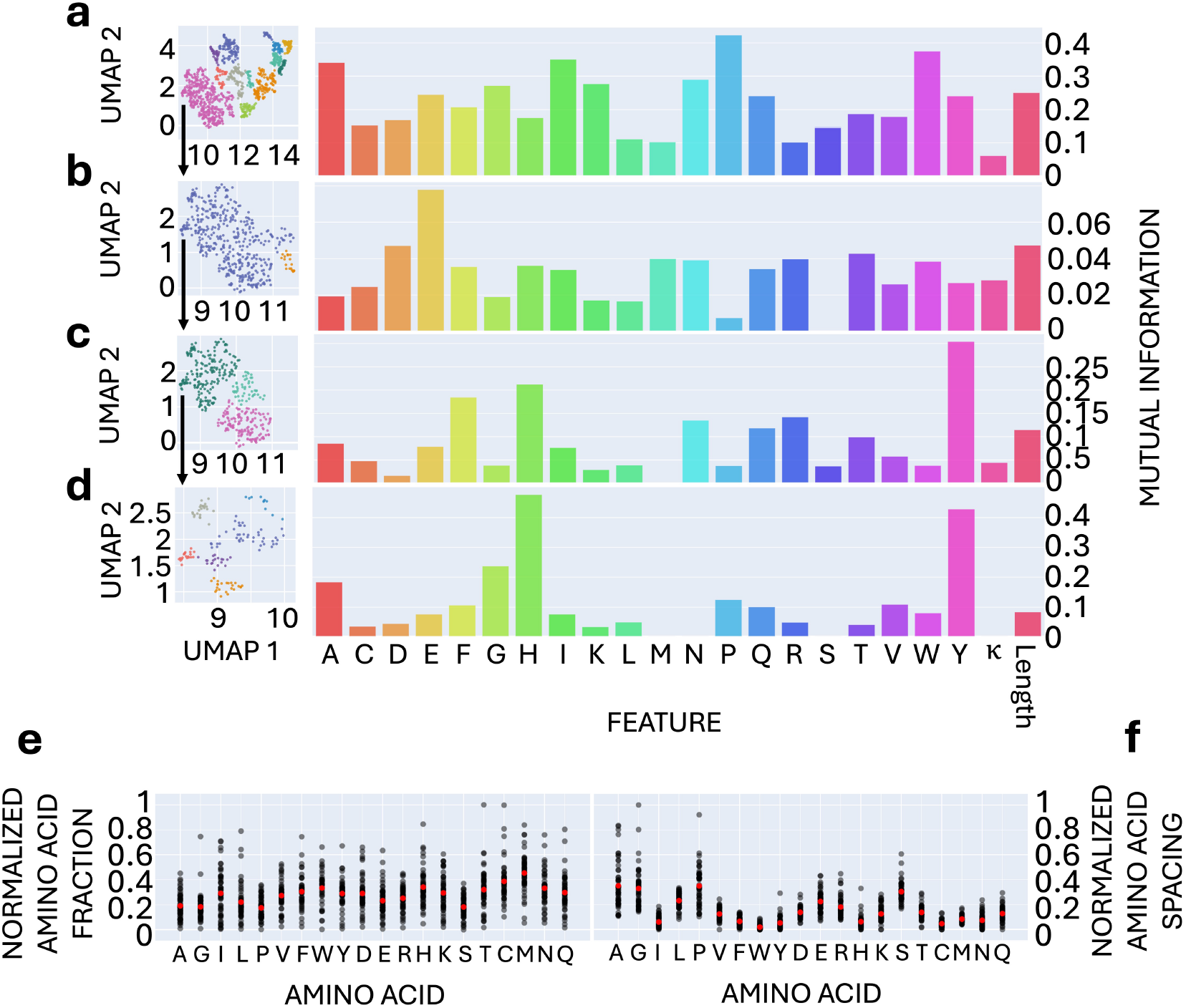
Clustering of the 1,529 elements in PR2 with its corresponding mutual information plot. a) Initial set splitting into 12 clusters. b) Subsequent clustering was applied only to the largest cluster (472 sequences). The first three rounds of subsequent clustering are shown in b) to d). Further clustering steps are excluded for visual clarity. Noise points are not included in the projections. Normalized averages per amino acid fraction (e) and identical amino acid spacing (f) for the obtained clusters (*black*) and global averages (*red*).

Further amino acid composition and spacing analyses (Figures 5f and 5g; 6e and 6f) revealed contrasting enrichment trends between PR1 and PR2: PR1 clusters were enriched in alanine, glycine, proline, leucine, glutamic acid, and serine; and depleted in aromatic residues and other hydrophobic amino acids, suggesting roles in flexible scaffolding and coactivator recruitment. PR2 clusters, by contrast, were distinctly enriched in hydrophobic amino acids, including phenylalanine and tryptophan, residues known to enhance activation^41^, and showed low conservation across most amino acids.

Importantly, the spacing distributions of the amino acids that dominate PR2 are very narrow, whereas the spacing distributions of amino acids within PR1 are very wide, i.e., none of the same type amino acids located particularly closely or in a repeating pattern. This is important because these closely located amino acids within PR2 can make sticker-like patches within the sequences that can form weak multivalent interactions. This suggests possibility of such patches within N-terminal portions of HTFs.

Full-sequence (FS) clustering (Figure S10) largely mirrored PR1 behavior, indicating that C-terminal effector segments dominate full-length compositional behavior.

### 0.6 Applying FALK22 to known activation and repression domains

To test whether FALK22 captures functional polarity within EDs, we projected the curated dataset of activation and repression domains^37^ into the FALK22-based reduced feature space. A clear gradient emerged from highly disordered (activation-dominated) to more ordered (repression-dominated) regions (Figure 7a and 7b). Strikingly, the effector domains from C2H2 ZF and NR family TFs, once again, clustered together near the lower-DS end (Figure 7c). This pattern reinforces our major finding that EDs and DBDs possibly co-evolved together, instead of evolving as modular entities. This pattern also indicates that FALK22 can separate EDs not only by DBD lineage but also by functional tendency toward activation or repression, reinforcing that disorder and composition jointly encode regulatory polarity.

**Figure 7.**
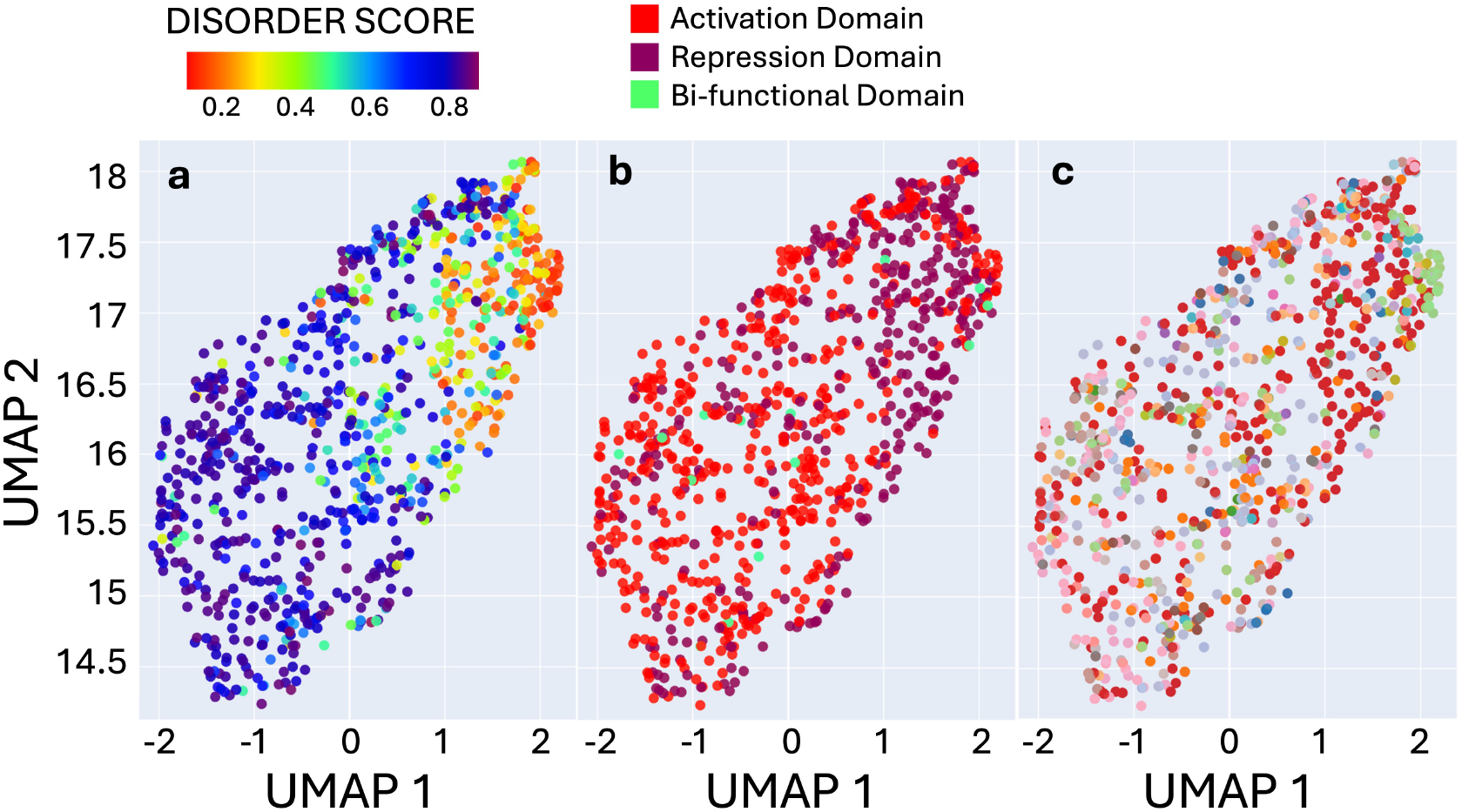
UMAP projections of the FALK22-processed known activation, repression and bi-functional domains (curated by Soto et al.^37^) The UMAP projections are colored according to a) DS, b) domain type [activation domain (*red*), repression domain (*purple*) and bi-functional domain (*green*)], and c) DBD family [follows the same color code as in Figure 2q].

## DISCUSSION

The systematic classification of HTFs based on their EDs represents a significant methodological advance that opens new frontiers in understanding transcriptional regulation. The first novelty in this work is to approximate EDs as non-DBDs of HTFs. This is significant because transcriptional activation or repression is a complicated process that does not follow a singular mechanism. Our vision here was based on the fact that any part of the TF chain that does not bind to DNA would be available for further interactions that can facilitate activation or repression.

Then the FALK22 framework that we have developed in this worksuccessfully revealed previously unrecognized patterns in amino acid composition and charge distribution that underlie the functional diversity of effector domains, establishing a foundation that bridges computational classification with emerging therapeutic paradigms and mechanistic insights. The identification of large number of distinct classes for PR1 and PR2, respectively, demonstrates remarkable functional diversity within HTFs.

The comparable performance of FALK22 compared to ESM embeddings in motif segregation and domain classification is particularly striking, given the computational simplicity of our biophysically-informed feature space relative to transformer-based approaches. This finding challenges the prevailing notion that complex deep learning models invariably outperform simpler, interpretable approaches in biological sequence analysis. The degradation of ESM performance with increasing model size—contrary to expectations based on structure prediction benchmarks—reveals important limitations in applying protein language models to sequence fragments and disordered regions. This observation has significant implications for the field, as it suggests that domain-specific approaches like FALK22 may be more appropriate for analyzing intrinsically disordered proteins than general-purpose language models trained on complete protein sequences.

Our analysis reveals fundamental differences between PR1 (C-terminal) and PR2 (mostly N-terminal) effector domains that provide crucial insights into transcription factor evolution and function. The conservation of amino acid composition in PR1 regions across diverse DBD families, coupled with enrichment in proline, leucine, glutamic acid, and serine, suggests these domains have evolved specialized roles in stabilizing transcriptional machinery and mediating promiscuous protein-protein interactions. This compositional signature aligns with the emerging understanding that C-terminal effector domains may serve as universal platforms for recruiting coactivators and affect transcription. The high compositional diversity observed in PR2 regions, with their enrichment in aromatic residues like phenylalanine and tryptophan, points to evolutionary pressure to develop context-specific activation mechanisms, potentially through *π*-*π* stacking interactions and unique binding interfaces that confer target gene specificity. The narrow distribution of amino acids enriched in PR2 suggests the presence of sticker-like patches in PR2 that promote, for example, condensate formation.

Our findings indeed provide crucial insights into the rapidly expanding field of transcriptional condensate biology. The charge patterning features captured by the *κ* parameter in FALK22, combined with the amino acid compositional differences we identified, directly relate to the phase separation propensities that drive biomolecular condensate formation. The prevalence of disorder-promoting residues and specific charge distributions in both PR1 and PR2 regions suggests that many human transcription factors possess the biophysical properties necessary for liquid-liquid phase separation. This connection is particularly relevant given recent advances demonstrating how transcriptional condensates contribute to gene regulation through the selective partitioning of transcription factors and coactivators.^76^ Future studies integrating our classification system with condensate biology could reveal how different effector domain classes contribute to the formation, composition, and function of transcriptional hubs.

Importantly, ED-like domains approximated as PR1 and PR2, showed DBD family-specific manifold gradients, e.g., C2H2 ZF and NR separated themselves from others consistently. This is striking because PR1 or PR2 databases completely excludes DBDs. No information regarding DBDs was fed to FALK22. Yet regardless of the feature identification technique (whether it is token embeddings from ESM or simple features from FALK22), the ED domain populated gradients segregated based on DBD families, which were also correlated with disorder score. This finding suggests co-evolution EDs with their DBD counterparts, aligning with their possibly shared regulatory logic.

The classification framework we present has also significant implications for the development of next-generation therapeutics targeting transcriptional dysregulation. The identification of distinct effector domain classes provides a roadmap for developing selective inhibitors that target specific transcription factor subfamilies. This precision approach could overcome the longstanding challenges associated with targeting the traditionally “undruggable” intrinsically disordered regions of HTFs. Recent breakthroughs in targeting intrinsically disordered proteins, exemplified by clinical compounds like ralaniten for androgen receptor effector domains and emerging condensate-modifying drugs (c-mods), demonstrate the therapeutic potential of this approach.^77,78^ Our classification system could accelerate similar drug discovery efforts by identifying effector domains with similar biophysical properties and suggesting common druggable features across HTF families.

The development of novel approaches for targeting intrinsically disordered regions, including the recently described “logos” system that uses AI to design binders for flexible protein regions,^79^ provides a technological foundation for translating our classification insights into therapeutic applications. The compositional and charge distribution patterns revealed by FALK22 could guide the design of such binders, enabling the development of highly selective transcription factor modulators. The integration of our classification system with high-resolution structural studies, such as cryo-electro microscopy and NMR spectroscopy represents a critical next step for understanding the mechanistic basis of the compositional patterns we identified.

Multi-omics integration approaches that combine our sequence-based classifications with transcriptomic, proteomic, and metabolomic data could provide systems-level insights into how effector domain diversity contributes to cell-type-specific gene expression programs. Such integrative analyses could reveal the regulatory logic underlying cellular identity and differentiation programs.

As the field moves toward more integrative and mechanistic approaches to studying gene regulation, the foundation provided by our effector domain classification system will serve as a valuable resource for the research community. The simple yet powerful framework we have established bridges the gap between sequence-level features and systems-level function, providing a roadmap for future investigations into the complex world of transcriptional regulation.

## Limitations of the study

While our classification framework represents a significant advance in understanding effector domain diversity, several important limitations highlight critical directions for future research and methodological development that will be essential for realizing the full potential of this approach. The reliance on existing database annotations for defining DBD boundaries introduces inherent uncertainties in effector domain delineation. The fact that only 1,588 of 1,632 HTFs have sufficient DBD annotations reflects a fundamental challenge in the field, i.e., the need for comprehensive experimental characterization of HTF domains. This limitation is particularly significant given that 57% of HTFs contain multiple DBDs, creating complex domain arrangements that our proxy region approach may not fully capture.

The exclusion of peptide fragments below 10 amino acids, while justified by secondary structure formation thresholds, may overlook functionally important short linear motifs that contribute to transcriptional regulation. Recent advances in understanding the role of short disordered regions in protein-protein interactions and phase separation suggest that future iterations of our approach should incorporate methods for analyzing these shorter sequences, potentially through specialized algorithms designed for motif discovery in disordered regions. Additionally, our feature space, while comprehensive in terms of amino acid composition, lacks information about higher-order sequence patterns and long-range correlations that may be important for effector domain function. Furthermore, many effector domains undergo post-translational modifications that can dramatically alter their functional properties, and our static sequence-based approach may not account for these dynamic changes. The role of cofactors, chromatin context, cellular localization, and tissue-specific expression patterns in modulating effector domain function cannot be captured by our current approach. Our methodology offers the opportunity to study the temporal and spatial resolution needed to describe the dynamics of gene expression regulation including the formation of phase-separated condensates,^80^ signaling,^81^ ligand binding,^82^ post-translational modifications (PTMs),^83,84^ homo-, heterodimerization,^85^ and nuclear transport^86^ which mainly take place across the ED. Future studies integrating proteomic data on post-translational modifications and cellular environment could significantly enhance the biological relevance of our classification system.

By acknowledging these limitations and embracing the opportunities they represent, the scientific community can build upon our foundational work to develop increasingly sophisticated and biologically relevant models of transcriptional regulation. The future of the field lies in integrative approaches that combine computational prediction, experimental validation, and systems-level understanding to unlock the full complexity of gene regulation in health and disease. Our classification framework provides a crucial starting point for this endeavor, offering a bridge between sequence-level features and functional outcomes that can guide future investigations and therapeutic developments.

## Acknowledgments

GHZ is a Cancer Prevention and Research Institute of Texas (CPRIT) scholar in cancer research and supported by CPRIT-RR220008, the Welch Foundation (Award E-2221 and Catalyst Center for Advanced Bioactive Materials Crystallization Award V-E-0001), and NSF CBET-2442006 (CAREER). The simulations presented in this work were performed using the computational resources provided by the Hewlett-Packard Enterprise Data Science Institute at the University of Houston. The authors thank Melissa Unlu, Preethi Kakarla, and Sandeep Reddy Kukunuru for their contributions in curating the HTFs datasets, and Heng Ma and the Argonne National Laboratory facilities for obtaining the ESM embeddings.

## Author contributions

Conceptualization, G.H.Z.; methodology, E.A., A.G., and A.R.; investigation, E.A., A.G., and N.S.; writing – original draft, E.A., A.G., and G.H.Z.; writing – review & editing, E.A., A.G., and A.R.; funding acquisition, G.H.Z.; resources, G.H.Z. and A.R.; supervision, G.H.Z.

## STAR METHODS

### Method details

#### 1.0.1 Datasets

The dataset of HTFs was extracted from the work of Lambert *et al*.^46^. We constructed a comprehensive dataset of 1,632 human transcription factors (TFs) and their corresponding 1,885 isoforms from the UniProtKB 2024 − 2 release^43^ along with their annotations. To identify the effector domains (ED), we removed the DBD segments from the main isoform sequence (identified throughout this work as full sequence (FS)). Since many TFs have multiple DBDs and DBDs are often located in non-terminal part of the sequence, the EDs approximated by simple removal of DBDs would be discontinuous, and therefore, can’t be used for classification. Hence, we identify all the continuous fragments of after the removal of known DBDs and consider the two longest of them, which we refer to as Proxy Region 1 (PR1) and Proxy Region (PR2), as representatives of the ED.

PR1 consisted only of the C-terminal sequences, while PR2 could be either the N-terminal segment or the longest sequence sandwiched between DBD sequences, only if it is at least 50 residues longer than the N-terminal fragment (Figure 1e). This process yielded sets of 1,588 EDs, 1,584 PR1s, and 1,566 PR2s. We classified the PR1 and PR2 sequences separately.

#### 1.0.2 Feature space

The sequence properties *κ* and disorder score (DS) were calculated using CIDER^87^ and Metapredict^56^, respectively. Normalized and averaged hydropathy score of chains were calculated using the per-amino acids scores assigned by Kyte and Dolittle^55^. All other parameters were calculated at physiological pH (7.4) using Biopython.^88^ The parameters *f* ^+^, *f* ^−^ and |*f* ^+^− *f* ^−^| are the fraction of basic amino acids, acidic amino acids, and mean net charge (MNC) (also known as absolute net charge per residue (NCPR)), respectively. After optimization of different feature spaces, we hypothesized that a total of 22 physically-interpretable features i.e., amino acid fractions, sequence length, and *κ* can sufficiently differentiate and classify the different sequences for the classification task.^89-91^

Because of the high dimensionality of the feature space, we implemented feature scaling and subsequent dimensionality reduction across the feature dataset to enhance computational efficiency during the clustering phase. We employed Uniform Manifold Approximation and Projection (UMAP)^92^ for this purpose, which is known for preserving global, local, and hierarchical structures within the data, thereby maintaining the relationships between data points. The optimal number of UMAP components was determined by evaluating the ‘trustworthiness’ score function available in `scikit-learn`^93^, where we iterated the number of UMAP components from 2 to 10. The selection was based on the first plateau observed in the plot of trustworthiness versus the number of UMAP components. Before applying UMAP, the physical features were normalized using the StandardScaler function from `scikit-learn` to ensure uniform scaling.

As an alternative feature representation, we also used token embeddings from the Evolutionary Scale Model (ESM)^42^, which is a fully contextualized language model pre-trained on the pre-clustered UniRef proteins datasets. The ESM embeddings were obtained from the last layer representation using different model sizes varying from 8M to 3B parameters. We applied zero padding to the sequences and mean pooling to obtain the final embeddings. Representation comparisons accross the manuscript were done against the 150M model size.

#### 1.0.3 Clustering

Within the clustering algorithms, density-based algorithms offer the advantage of identifying arbitrary-shaped clusters in the data, without requiring the user to define the expected number of clusters beforehand.^94^ We applied Hierarchical Density-Based Spatial Clustering of Applications with Noise (HDBSCAN) which can identify non-linear relations between data points and segregate them based on the hyperparameters.^57^

Among HDBSCAN’s tunable parameters, the minimum cluster size (MCS) and minimum sample size (MSS) exert the greatest influence on clustering outcomes. We systematically varied both parameters from 2 to 30 (Figure S9) and assessed the resulting clusterings using two complementary criteria: (i) the Density-Based Clustering Validation (DBCV) score,^95^ which indicates the separation of the resulting clusters (higher values indicate better-defined clusters), and (ii) the noise fraction, representing the proportion of sequences not assigned to any cluster.

While maximizing DBCV score is desirable, the highest scores often coincided with trivial outcomes, such as single dense cluster or solutions dominated by noise, resulting in poor biological interpretability. Therefore, we selected the parameter combination that achieved a balanced trade-off: a moderate DBCV score, an acceptable noise fraction, and a meaningful number of clusters.

To further homogenize the resulting clusters, we performed successive independent HDB-SCAN clustering to any cluster containing more than 100 sequences. Reclustering continued until each subcluster contained fewer than 100 sequences or the overall noise fraction surpassed 0.3. At every step, we computed the mutual information between cluster identity and individual features using the `mutual info classif` function of `scikit-learn`^93^. The resulting mutual-information values quantify how strongly each feature contributes to the classification; higher values indicate greater influence on cluster assignment.

## Additional resources

Cell.com homepage: https://www.cell.com

Templates for Cell Press authors: https://www.cell.com/templates

