## Supporting Figures and Tables for "Classification of Human Transcription Factors Based on Their Effector Domains via Unsupervised Learning"

### Supporting information for "Classification of Human Transcription Factors Based on Their Effector Domains via Unsupervised Learning"

Eduardo Ayala[1,4], Ayush Gupta[1,4], Nehil Shreyash[1], Arvind Ramanathan[2], and Gül H. Zerze[1,3,4,5,6,*]

[1]William A. Brookshire Department of Chemical and Biomolecular Engineering, University of Houston, Houston, TX, USA
[2]Argonne National Laboratory, Lemont, IL, USA
[3]Present address: 4226 Martin Luther King Boulevard Houston, TX, USA
[4]These authors contributed equally
[5]Senior author
[6]Lead contact

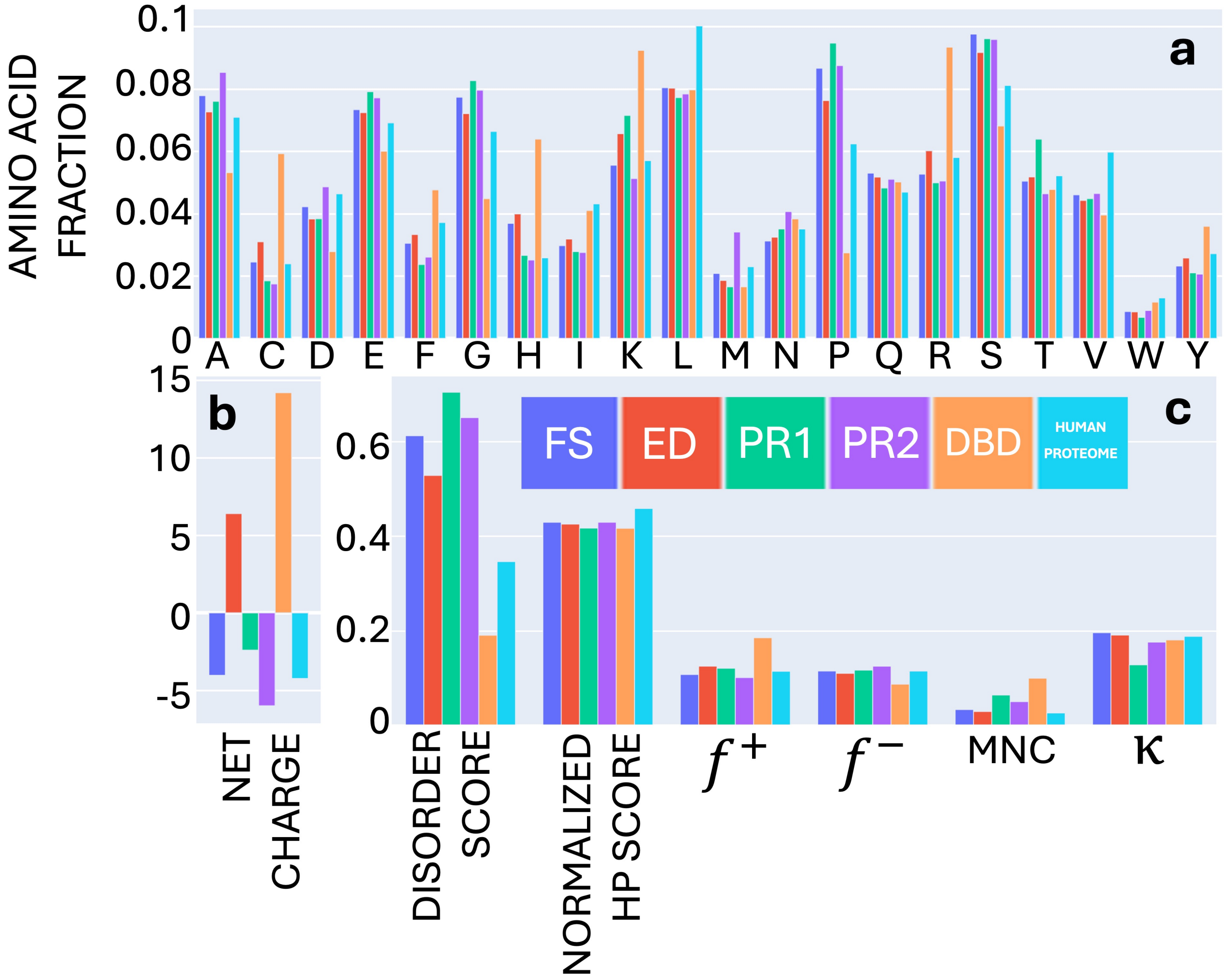


**Figure S1. Amino acid fraction and physical properties averages comparison of the Full Fequence (FS), Effector Domains (ED), DNA Binding Domains (DBD), Proxy region 1 (PR1), Proxy region 2 (PR2), and Human Proteome datasets.**

a) Amino acid fraction composition, b) protein net charge, and c) disorder score (DS), normalized hydropathy score, positive net charge ($f+$), negative net charge ($f-$), mean net charge (MNC), and $\kappa$.

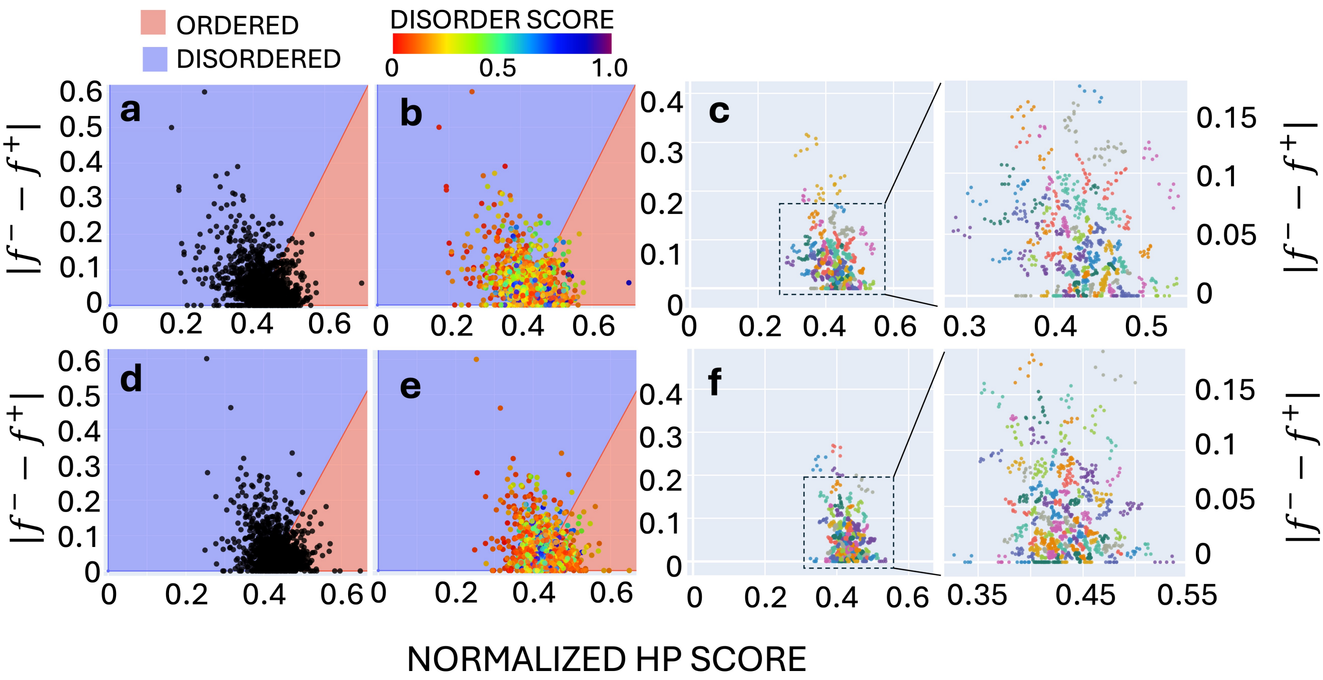


**Figure S2. Uversky plot, $|f^+ - f^-|$ versus normalized hydropathy score.**

Scatter plots for a) PR1 and d) PR2 in ordered $(red)$ and disordered $(blue)$ protein regions. DS distribution for PR1 (b) and PR2 (e). HDBSCAN clustering results over the $|f^+ - f^-|$ versus normalized hydropathy score space for c) PR1 and f) PR2. Sequences classified as noise are not shown for clarity in the resulting clusters.

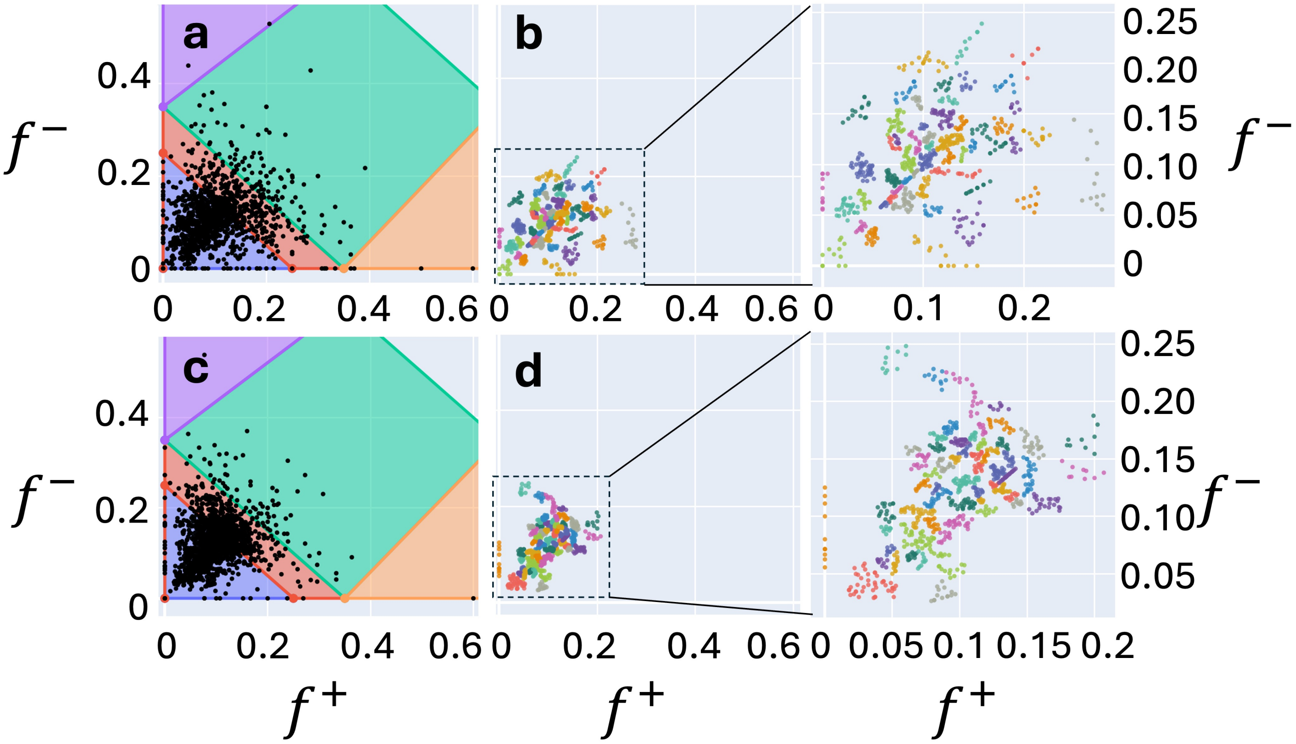


**Figure S3. Das Pappu diagram of state $f^+$ versus $f^-$ for PR1 (a) and PR2 (c).**

R1 ($blue$): weak polyampholytes or weak polyelectrolytes that form globules or tadpole-like conformations. R3 ($green$): strong polyampholytes that form non-globular conformations, such as coil-like, hairpin-like, or a mixture. R2 ($red$), continuum of conformations between those in R1 and R3. R4 ($green$ and $orange$): strong polyelectrolytes which sample coil-like conformations approaching the excluded-volume limit. Clustering results over the $f^+$ versus $f^-$ space for PR1 (b) and PR2 (d). Sequences classified as noise are not shown for clarity in the resulting clusters.

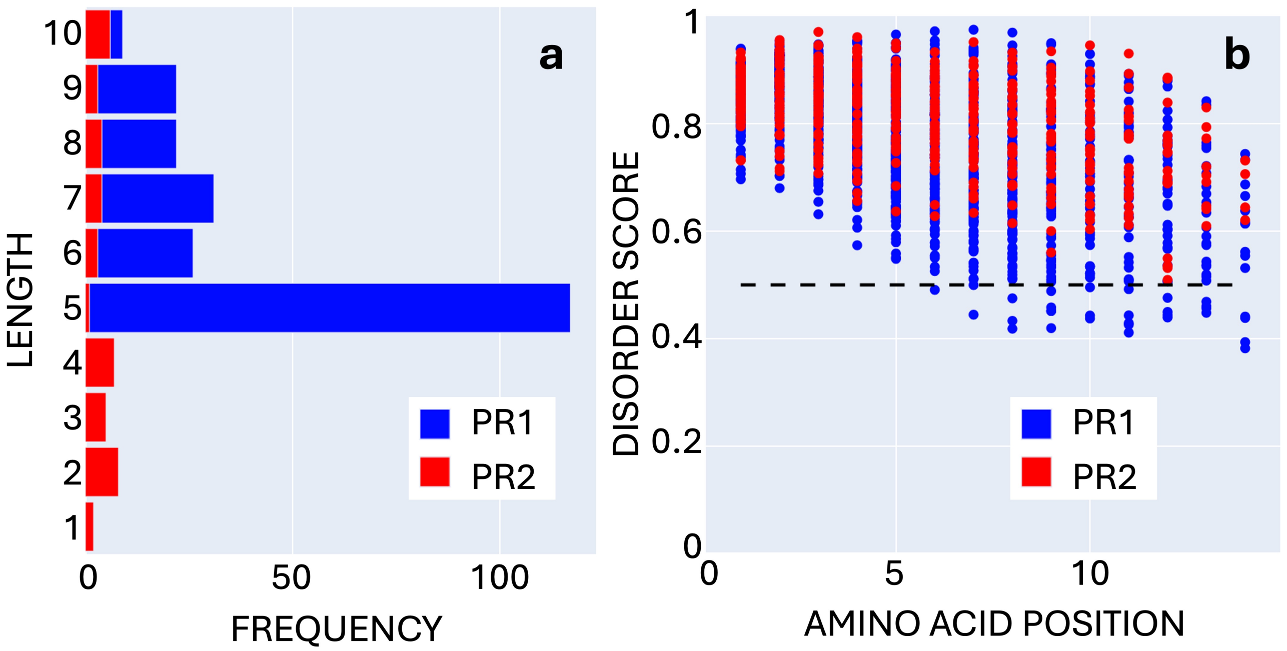


**Figure S4. Size distribution and per-residue DS for effector domains under 10 aa length.**

Fragment description of short effector domains PR1 (blue) and PR2 (red): a) Length distribution, and b) Per-residue DS within the chain.

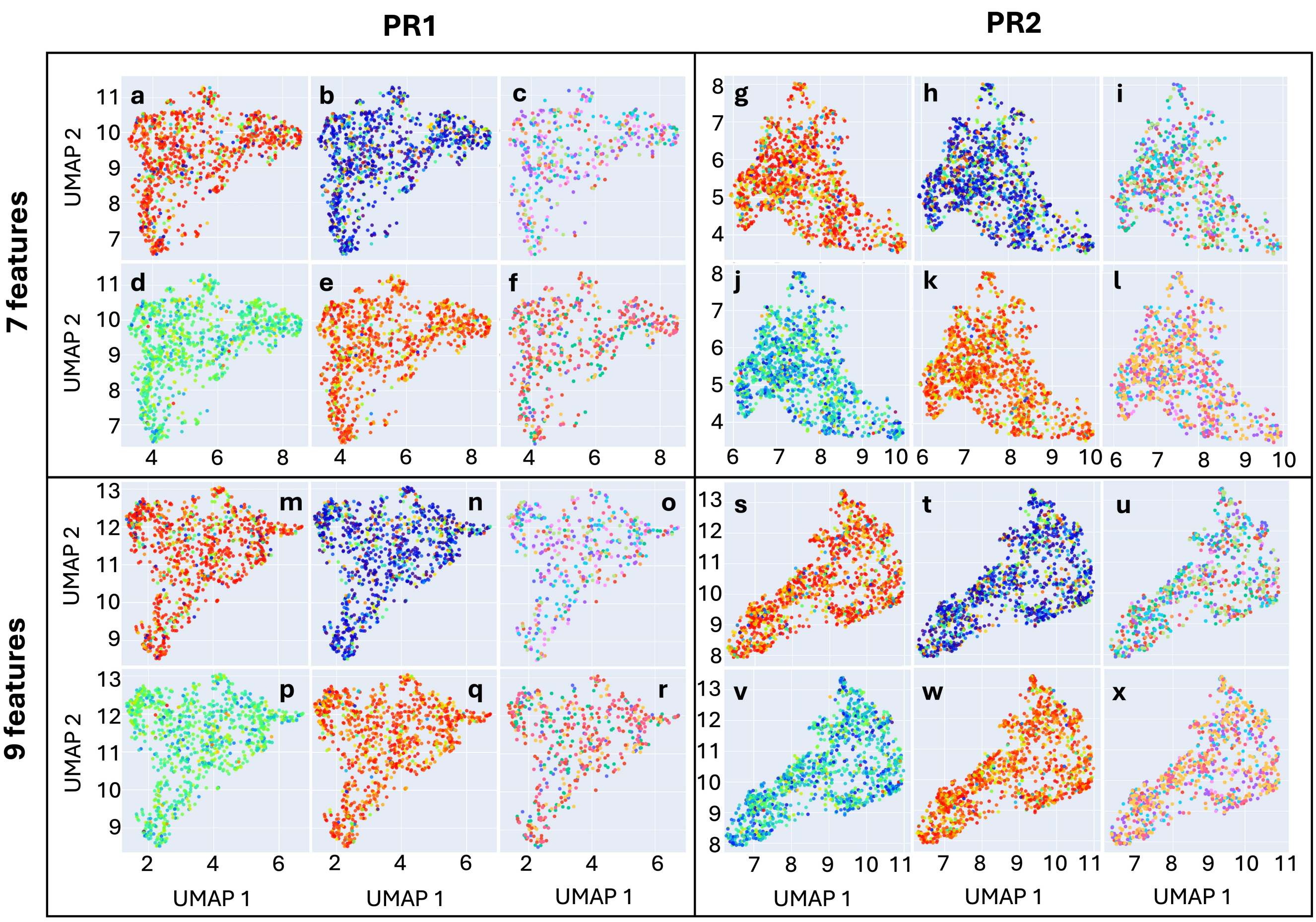


### Figure S5. Two-dimensional projection of the feature spaces F7 and F9 for PR1 and PR2.

Seven-feature space for PR1 (a - f), PR2 (g - l) and nine-feature space for PR1 (m - r) and PR2 (s - x). Each feature space is colored using the same order of scales: Family color code (a, g, m, s), Average DS (b, h, n, t), FIMO motifs (c, i, o, u), mean hydrophobicity (d, j, p, v), mean net charge (e, k, q, w), and STREME motifs (f, l, r, x).

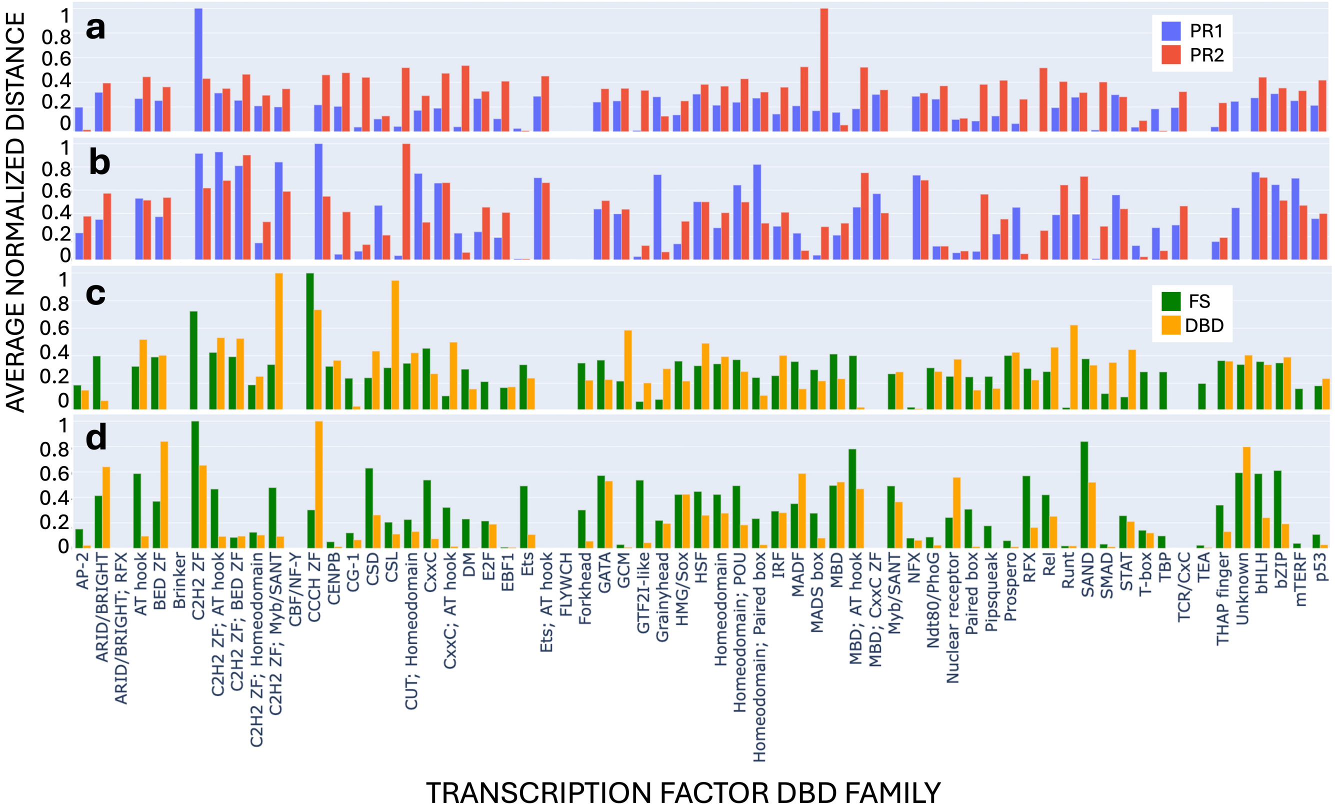


#### Figure S6. Average distance between members of the same DBD family on two-dimensional projections (from Figure 2).

Normalized Euclidean distance between sequences within the same DBD family for PR1 ($blue$) and PR2 ($red$) from the two-dimensional projections of Figure 2 for FALK22 (a) and ESM (b), and for FS ($green$) and DBD ($yellow$) from the two-dimensional projections of Figure 2 for FALK22 (c) and ESM (d).

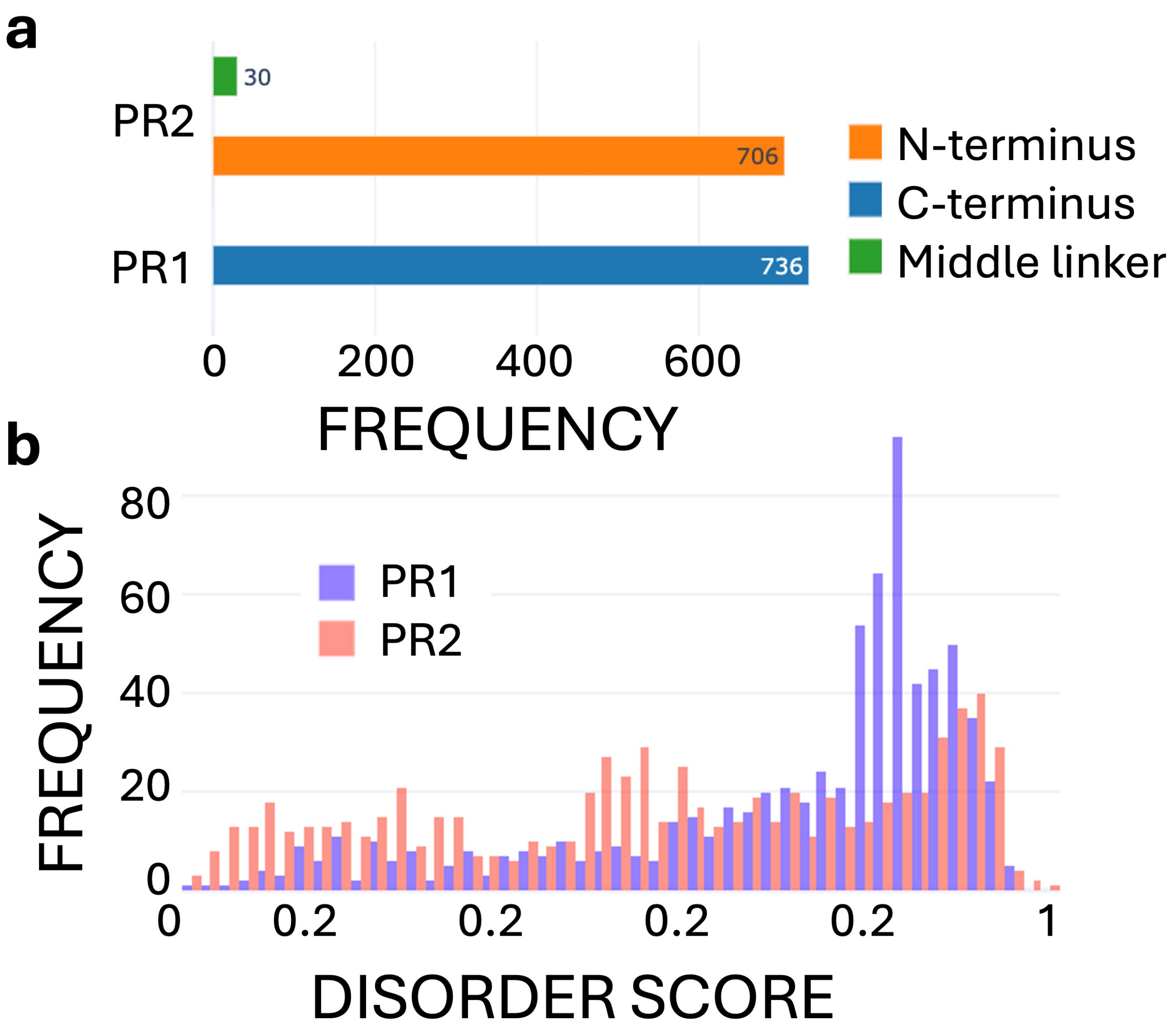


**Figure S7. Zinc finger proxy regions location and their DS distribution.**

a) The locations of PR1 and PR2 in zinc finger sequences. The C-termini are populated the PR1 dataset exclusively, while the N-terminus and longer middle linkers contribute to the PR1 dataset. Only 30 of the 736 ZF PR2's are middle linkers. b) DS distribution of PR1 and PR2 of ZFs.

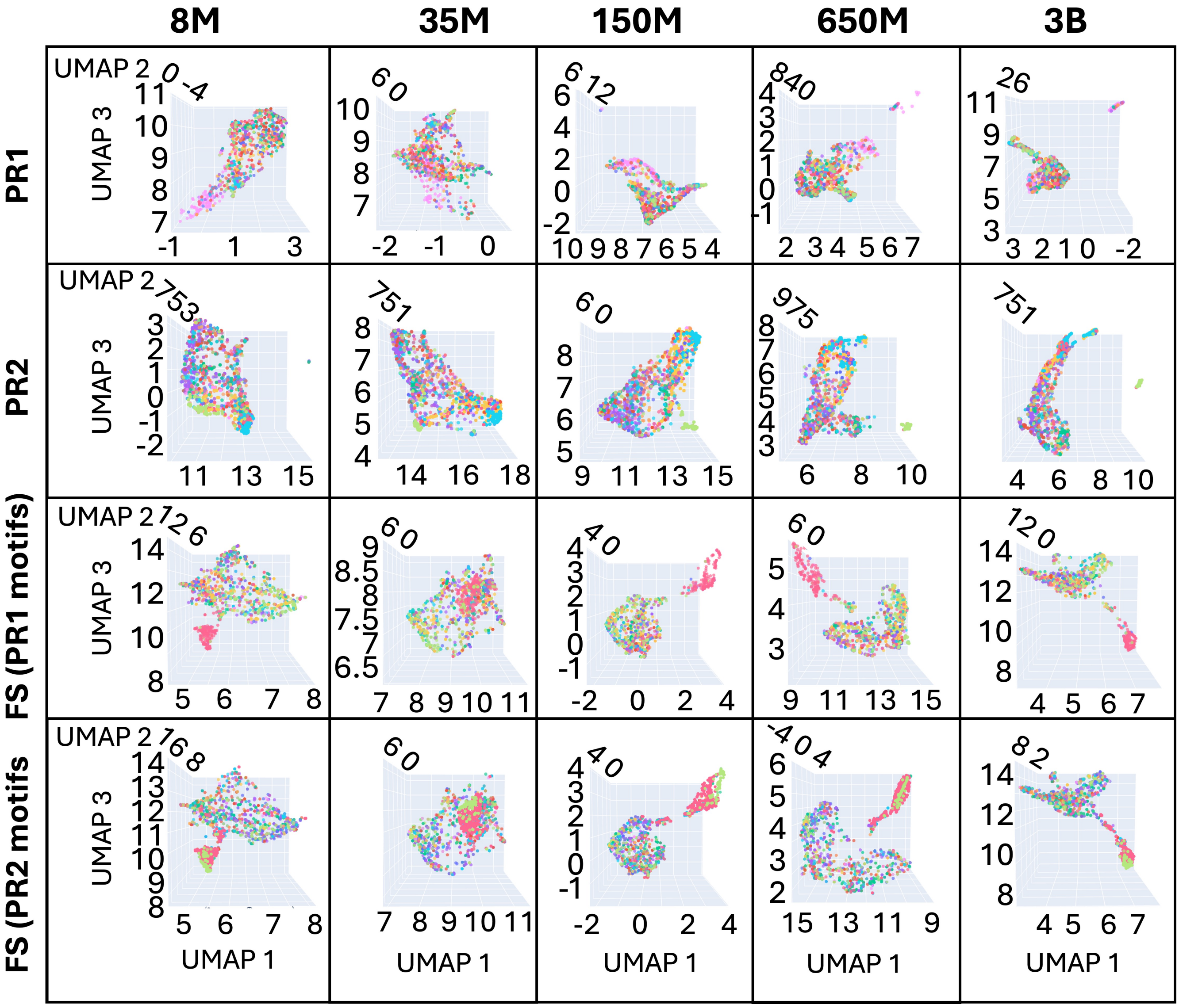


**Figure S8. Three-dimensional projections of the different ESM model size embeddings.**

Projection of ESM-based features (token embeddings) for PR1, PR2, and FS datasets colored by the STREME and FIMO motif search results for the ESM model sizes 8M, 35M, 150, 650M, and 3B parameters.

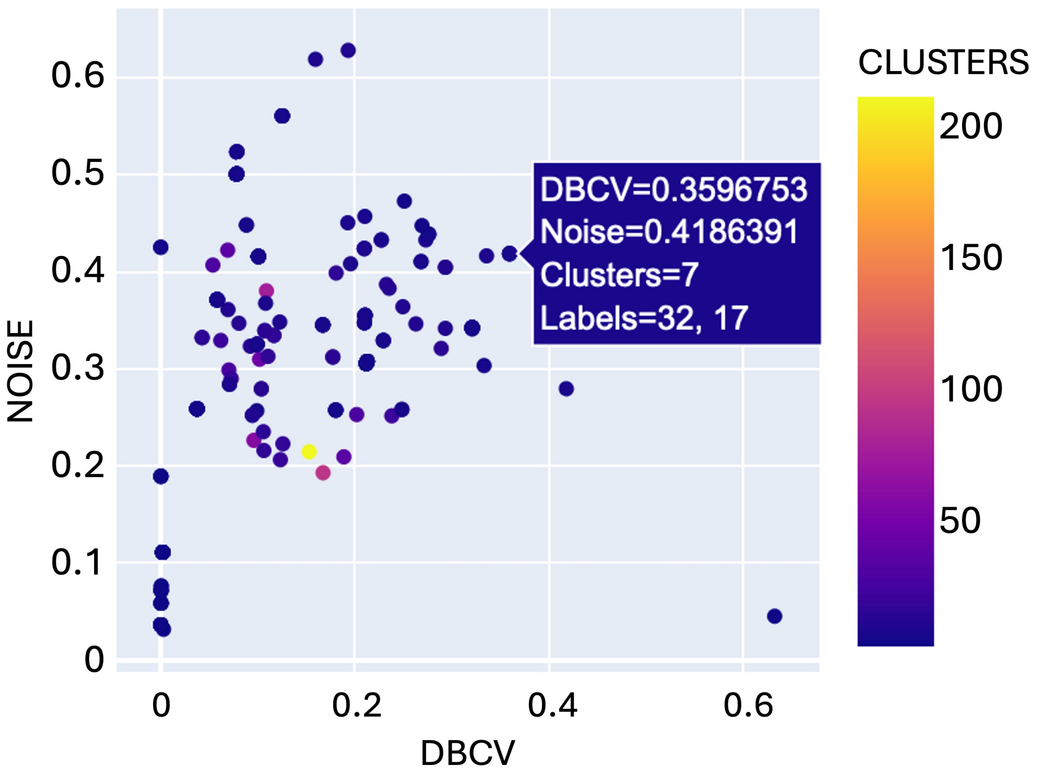


**Figure S9. Noise vs DBCV Score plot.**

Individual points indicate each HDBSCAN trial using a combination of hyperparameters MCS and MSS. The line 'Labels' on the legend indicates the MCS and MSS for the corresponding point.

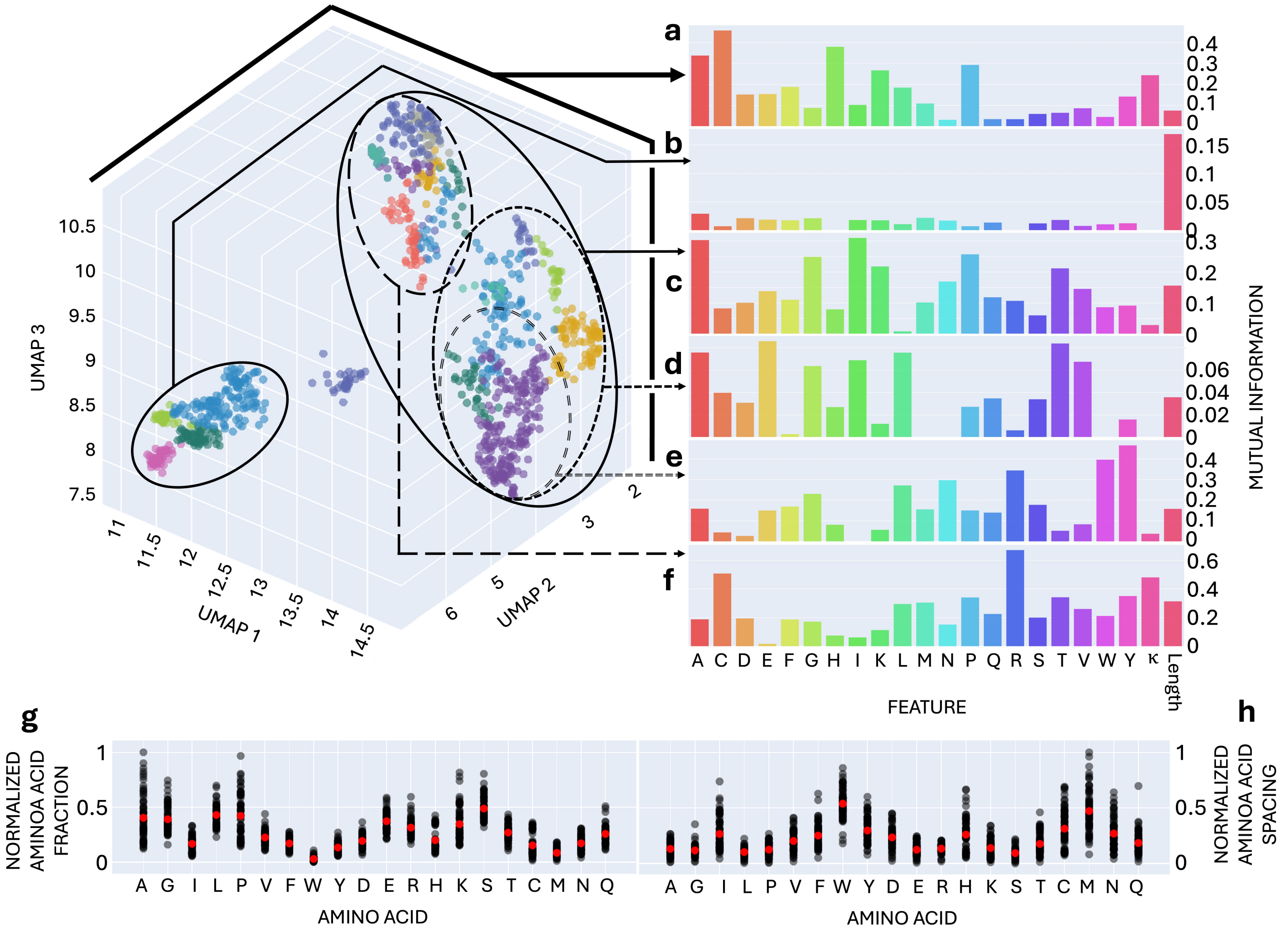


#### Figure S10. FS feature space, subsequent clustering and cluster analysis.

Clustering of the 1,632 elements in FS with its corresponding mutual information plot. b) Initial set splits into 3 clusters with 35, 371 and 1,202 sequences. Projections included for the subsequent clustering of the 371 elements (c) and the 1,202 elements cluster (a), 1,141 (d), 390 (e), and 309 (f). Noise points are not included in the projections. Normalized averages per amino acid fraction (g) and identical amino acid spacing (h) for the obtained clusters ($black$) and global averages ($red$).

| DBD-family | PR1 | | PR2 | | FS | |
|---|---|---|---|---|---|---|
| | FALK22 | ESM | FALK22 | ESM | FALK22 | ESM |
| Nuclear Receptor | ■ | ■ | | | ■ | |
| C2H2 ZF | | | ■ | ■ | | |
| Homeodomain | | | | ■ | | |
| Forkhead | ■ | ■ | | | | |
| Homeodomain; POU | | | | ■ | | ■ |
| Homeodomain; Paired Box | ■ | ■ | | | | |
| T-box | ■ | ■ | | ■ | | ■ |
| CSD | | ■ | | | | ■ |
| STAT | | ■ | ■ | ■ | ■ | ■ |
| Paired Box | ■ | ■ | | | | |
| DM | | ■ | | | | |
| Grainyhead | | ■ | | | ■ | ■ |
| SMAD | ■ | ■ | | | ■ | ■ |
| MBD | | ■ | | | | |
| HMG / Sox | | | | ■ | | |
| IRF | | ■ | | | | |
| CENPB | ■ | | | | ■ | ■ |
| Myb / SANT | | | | | ■ | |
| E2F | | | | | ■ | |
| MADS Box | ■ | | | | | |
| RFX | ■ | | | | | |
| TEA | ■ | | | | | |
| BED ZF | ■ | | | | | |

**Table S1. Summary of major DBD families exhibiting spatial clustering in the reduced feature spaces derived from FALK22 and ESM embeddings for PR1, PR2, and full-length *FS) datasets.**

Families that form compact clusters in the two-dimensional projections are marked with green boxes, indicating spatial proximity of sequences within the same DBD family. Green-filled boxes therefore correspond to the presence of visually detectable clusters in the respective spaces.

**Table S2. RMSD Values after alignment of FALK22 and ESM (different model sizes) embeddings in the three-dimensional space.**

| ESM Model Size | 8M | 35M | 150M | 650M | 3B |
|---|---|---|---|---|---|
| FS | 0.3598 | 0.5954 | 0.3152 | 0.3028 | 0.5739 |
| PR1 | 0.4314 | 0.4924 | 0.6025 | 0.4572 | 0.4764 |
| PR2 | 0.4256 | 0.3928 | 0.3626 | 0.4276 | 0.5515 |
